# Local population structure and patterns of Western Hemisphere dispersal for *Coccidioides spp*., the fungal cause of Valley Fever

**DOI:** 10.1101/024778

**Authors:** David M. Engelthaler, Chandler C. Roe, Crystal M. Hepp, Marcus Teixeira, Elizabeth M. Driebe, James M. Schupp, Lalitha Gade, Victor Waddell, Kenneth Komatsu, Eduardo Arathoon, Heidi Logemann, George R. Thompson, Tom Chiller, Bridget Barker, Paul Keim, Anastastia P. Litvintseva

**Affiliations:** TGen North, Translational Genomics Research Institute, Flagstaff, AZ; Informatics and Computing Center, Northern Arizona University, Flagstaff, AZ; Mycotic Diseases Branch, National Center for Emerging and Zoonotic Infectious Diseases, Centers For Disease Control and Prevention, Atlanta, GA; Division of Public Health Services, Arizona Department of Health Services, Phoenix, Arizona; Asociación de Salud Integral, Guatemala City, Guatemala; Universidad de San Carlos, Ciudad Universitaria, Guatemala City, Guatemala; Division of Infectious Diseases, Department of Medicine, University of California Davis, Davis, CA; Microbial Genetics and Genomics Center, Northern Arizona University, Flagstaff, AZ

**Keywords:** *Coccidioides*, molecular epidemiology, phylogenetics, mycology

## Abstract

Coccidioidomycosis (or Valley Fever) is a fungal disease with high morbidity and mortality that affects tens of thousands of people each year. This infection is caused by two sibling species, *Coccidioides immitis* and *C. posadasii*, which are endemic to specific arid locales throughout the Western Hemisphere, particularly the desert southwest of the United States. Recent epidemiological and population genetic data suggest that the geographic range of coccidioidomycosis is expanding as new endemic clusters have been identified in the state of Washington, well outside of the established endemic range. The genetic mechanisms and epidemiological consequences of this expansion are unknown and require better understanding of the population structure and evolutionary history of these pathogens. Here we perform multiple phylogenetic inference and population genomics analyses of 68 new and 18 previously published genomes. The results provide evidence of substantial population structure in *C. posadasii* and demonstrate presence of distinct geographic clades in central and southern Arizona as well as dispersed populations in Texas, Mexico, South America and Central America. Although a smaller number of *C. immitis* strains were included in the analyses, some evidence of phylogeographic structure was also detected in this species, which has been historically limited to California and Baja Mexico. Bayesian analyses indicated that *C. posadasii* is the more ancient of the two species and that Arizona contains the most diverse subpopulations. We propose a southern Arizona-northern Mexico origin for *C. posadasii* and describe a pathway for dispersal and distribution out of this region.

## Importance

Coccidioidomycosis, or Valley Fever, is caused by the pathogenic fungi *Coccidioides posadasii* and *C. immitis*. The fungus and disease are primarily found in the American desert Southwest, with spotted distribution throughout the Western Hemisphere. Initial molecular studies suggested a likely anthropogenic movement of *C. posadasii* from North America to South America. Here we comparatively analyze eighty-six genomes of the two *Coccidioides* species and establish local and species-wide population structure to not only clarify the earlier dispersal hypothesis but also provide evidence of likely ancestral populations and patterns of dispersal for the known subpopulations of *C. posadasii*.

## Introduction

*C. immitis* and *C. posadasii* are the etiological agents of coccidioidomycosis, or Valley Fever, a primarily pulmonary disease that causes tremendous morbidity (i.e., thousands of new infections per year) in the Southwestern US and other focal regions in the Americas (1). *C. posadasii* was first characterized as “non-California *C. immitis*” found in the Southwestern US states of Arizona (AZ), New Mexico (NM), Texas (TX), Mexico and sporadic locales in Central and South America, and later named as a distinct species in 2002 (2). *C. immitis* is found primarily in California’s Central Valley and the southern California-Baja Mexico region and has been recently described as endemic in southeastern Washington state (3). The two species are similar by clinical and microbiological phenotype, although specific differences have been reported (4,5). *Coccidioides* is a dimorphic fungus, with a saprobic phase consisting of mycelia in the soil and a parasitic phase from inhaled arthroconidia, resulting in mammalian infection (6). Death and burial of the mammalian host can lead to re-infection of the soil (7). Both species of *Coccidioides* follow these stages and appear to lead to similar clinical outcomes (8).

Previous genomic analyses of targeted loci (e.g., microsatellites) led to the understanding that the California populations and the non-California populations are genetically distinct species (2,9). Whole genome analysis has supported this concept, with hundreds of thousands of single nucleotide polymorphisms (SNPs) separating the two species (10). In a comparative analysis of a single genome from each species, Sharpton et al (8) identified 1.1-1.5 Mb (~4-5%) of dissimilar genetic content between the two genomes. A later comparative genomics study, based on the analysis of ten genomes of each species, determined that the separation of the two species has not remained fully complete, and documented possible presence of introgression from hybridization events following species separation (11). As rapid DNA sequencing analysis has become more accessible to the research and public health laboratories, approaches like whole genome SNP typing (WGST) have been shown to be useful for identifying clonal fungal outbreaks (12–15). However, genomic epidemiology is also needed to help establish location of exposure of a non-clonal outbreak and robust phylogenomic analysis is needed for in-depth investigation of local population structure to better understand pathogen emergence, dispersal and expansion (15).

It is currently thought that wind, water, mammal hosts and anthropogenic causes are important mechanisms for local and geographic-scale dispersal of arthroconidia (16–17). Using microsatellite data from 163 isolates, Fisher *et al* (9) described a phylogeographic pattern of *C. posadasii* that was hypothesized to have been driven by human migration, strongly suggesting a movement of *Coccidioides* into South America that was contemporary with the first human movements onto the continent. While this groundbreaking study was critical to understanding large-scale phylogeographic features of *Coccidioides* epidemiology, and evidence was identified for local population structure, additional microsatellite analysis of Arizona-only isolates was unable to identify local structure within that state (18).

Given the limited number of *Coccidioides* genomes previously available and the complexities of fungal recombination (19), an accurate understanding of population structure and phylogeography has been problematic. Here we provide comparative phylogenomic and recombination analysis of 86 *Coccidioides* genomes with some insight into the population structure, mutation rates and phylogeographic patterns between and within the two species, as well as a proposed model for the global dispersal and distribution of *C. posadasii*.

## Results

### Genome sequencing and SNP calling

Sixty-eight newly sequenced genomes were compared to 18 of the existing 28 published genomes (11,12,14), resulting in a total of 86 genomes (26 *C. immitis* and 60 *C. posadasii)* available for analysis (Table S1). Depth of sequence coverage for the new genomes ranged from 23X to 228X (avg. 67X). The two assembled genomes (San Diego_1 and B10813_TX) used as references had a total of 28,389,157 and 28,389,157 bases, respectively. San Diego_1 assembled into 3,464 contigs with an n50 of 200,441 bases. B10813_TX assembled into 2,254 contigs and had an n50 of 43,956 bases. The publically available Sanger sequencing data (http://www.broad.mit.edu/annotation/genome/coccidioides_group/-MultiHome.html) only included final assemblies, making it difficult to distinguish true SNPs from sequence error; as such, these genomes were not included in all population analyses. Six of the previously published Illumina sequences (11) had low depth of coverage (3.4-8.5X) and the three previously published SOLiD sequences (12) used short reads (35-50bp) resulting in larger error rates (data not shown) and these sets were removed from the analyses. The genus level SNP analysis included the 69 genomes (18 *C. immitis* and 51 *C. posadasii*) with highest coverage and quality, excluding the previously published genomes to maximize the number of identifiable SNPs. That analysis identified 405,244 shared SNP loci, that is, loci in genetic content shared by all strains (Figure 1), with 296,632 of those being parsimony informative. A more inclusive phylogenetic analysis, using available Sanger sequenced isolates included 81 genomes (22 *C. immitis* and 59 *C. posadasii*) (Supplemental Figure 1), found only 128,871 shared SNP loci. The total number of SNPs was reduced likely due to filtering out of low quality SNPs in the previously published Sanger sequence data, however the overall topologies did not demonstrably change between the conservative and inclusive phylogenies.

**Figure 1.**
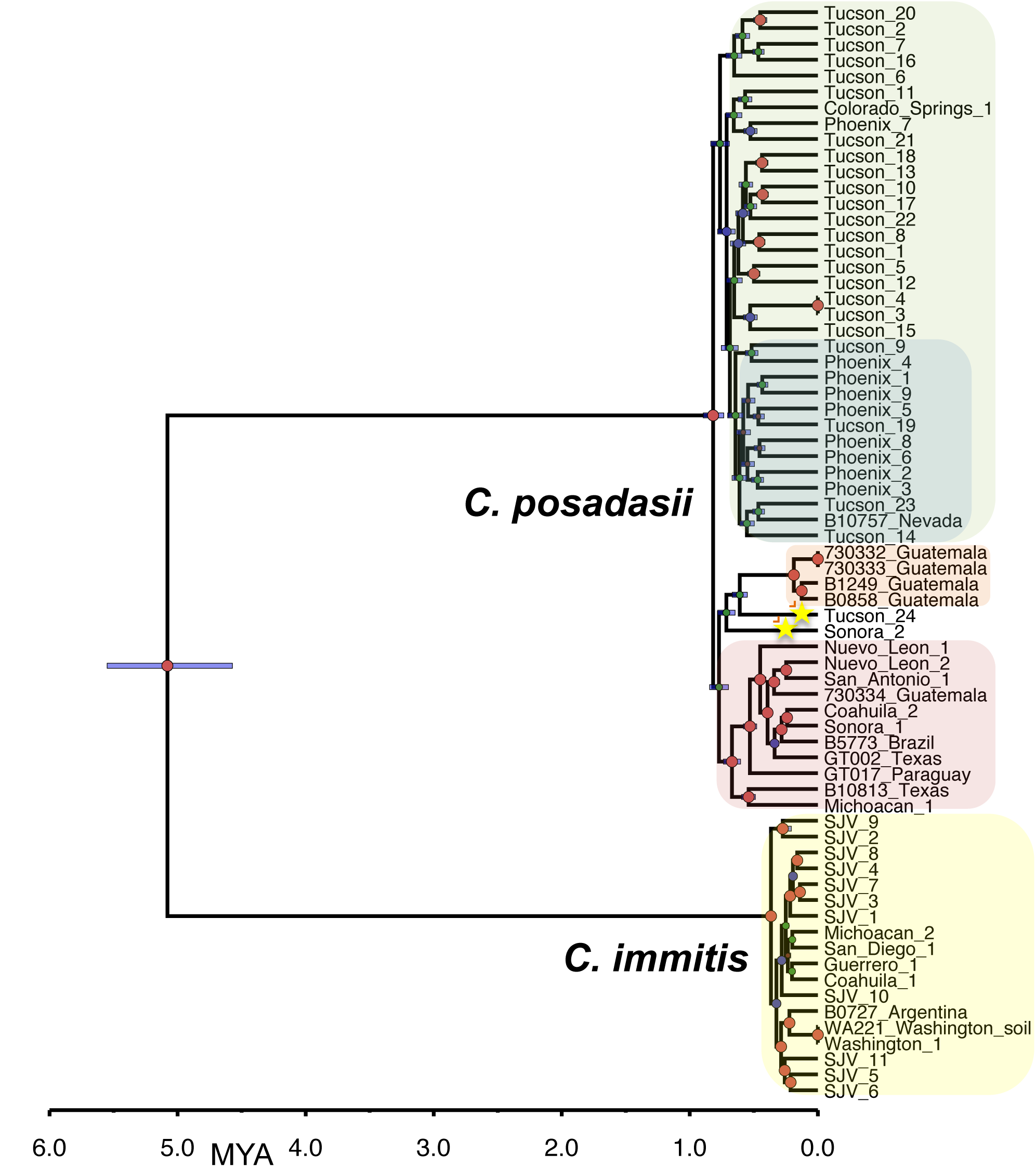
Bayesian phylogenetic analysis of *C. immitis* and *C. posadasii* isolates from all known endemic regions. The Bayesian statistical framework incorporated in BEAST 1.8.1 was used to integrate prior information, in the form of internal node timing estimates (from a fossil record) with a rooted tree, to produce a cahbrated phylogeny. The analysis was performed on WGS SNP data from 69 *Coccidioides* genomes, 18 *C. immitis* and 51 C. *posadasii*. Clades of interest are highlighted: green: Arizona, blue: Phoenix subclade of Arizona, orange: Guatemala, pink: Texas-Mexico-South America, and yellow: *C immitis*. Stars highlight strains of interest. Posterior probabilities are indicated as node size. Purple node bars are shown for each node and are informative for the 95% confidence interval for the timing estimate. The timeline represent millions of years before the present.

A large proportion of the parsimony informative SNPs (28,660; 40.7%) separate the two species (additional genus level Bayesian, likelihood, and parsimony trees can be found in Supplemental material). Additionally, an analysis of molecular variance (AMOVA) showed significantly high levels of genomic separation (ΦPT = 0.91, p<0.001) providing clear support for their taxonomical distinction. *C. posadasii* demonstrated a larger average SNP distance (i.e., the number of base substitutions per site from averaging overall pairwise distance between isolates within each species) than *C. immitis* (0.062 versus 0.024 respectively) (Table 1). Furthermore, Bayesian estimations of molecular rate suggest C. *posadasii* as having a much earlier within-species divergence than *C. immitis* (Figure 1 and below).

**Table 1.**
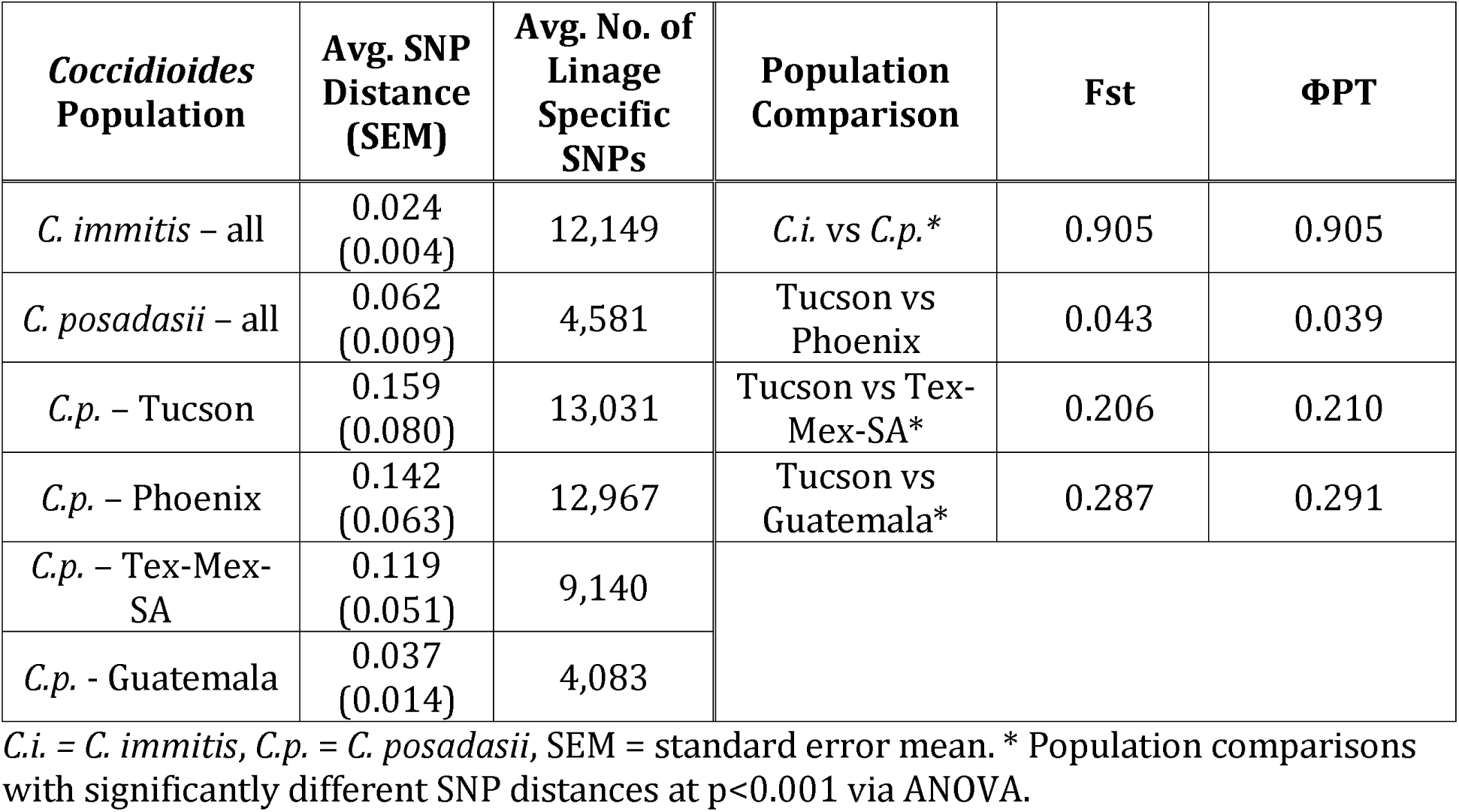
Summary statistics of *Coccidioides* population analyses.

| *Coccidioides*<br>Population | Avg. SNP<br>Distance<br>(SEM) | Avg. No. of<br>Lineage<br>Specific<br>SNPs | Population<br>Comparison | Fst | ΦPT |
| --- | --- | --- | --- | --- | --- |
| *C. immitis* – all | 0.024<br>(0.004) | 12,149 | *C.i.* vs *C.p.* * | 0.905 | 0.905 |
| *C. posadasii* – all | 0.062<br>(0.009) | 4,581 | Tucson vs<br>Phoenix | 0.043 | 0.039 |
| *C.p.* – Tucson | 0.159<br>(0.080) | 13,031 | Tucson vs Tex-Mex-SA* | 0.206 | 0.210 |
| *C.p.* – Phoenix | 0.142<br>(0.063) | 12,967 | Tucson vs<br>Guatemala* | 0.287 | 0.291 |
| *C.p.* – Tex-Mex-SA | 0.119<br>(0.051) | 9,140 |  |  |  |
| *C.p.* - Guatemala | 0.037<br>(0.014) | 4,083 |  |  |  |

*C.i.* = *C. immitis*, *C.p.* = *C. posadasii*, SEM = standard error mean. \* Population comparisons with significantly different SNP distances at $p < 0.001$ via ANOVA.

### Coccidioides posadasii population structure

Phylogenetic histories of each species were interrogated separately, using multiple phylogenetic inference and population genomics tools, as no one tool can provide a complete phylogenomic picture. The *C. posadasii* analysis was primarily based on 51 genomes: 38 isolates from the US, six isolates from Mexico, five from Central America (Guatemala) and three from South America (Brazil, Paraguay and Argentina). Isolate locales are based on available metadata and may represent location of sample isolation, laboratory that stored isolate, or home of patient (Supplemental Table 1). A total of 253,291 SNPs from shared loci (134,752 parsimony informative) were identified among the *C. posadasii* isolates. The pairwise homoplasy index test (phi) test found statistical evidence for recombination (p=0.0), however three major genetic populations were clearly separated and well supported by standard population statistics (Table 1). Evidence from three Bayesian analyses: a) BEAST (Figure 2); b) *fineStructure* (Figure 4); and BAPS (Figure 5), as well as the maximum parsimony and maximum likelihood (ML) bootstrap analyses (Supplemental Figures 3 and 4), all support this separation into three geographically-related populations: 1) Arizona; 2) Texas, Mexico and South America; and 3) Guatemala.

**Figure 2.**
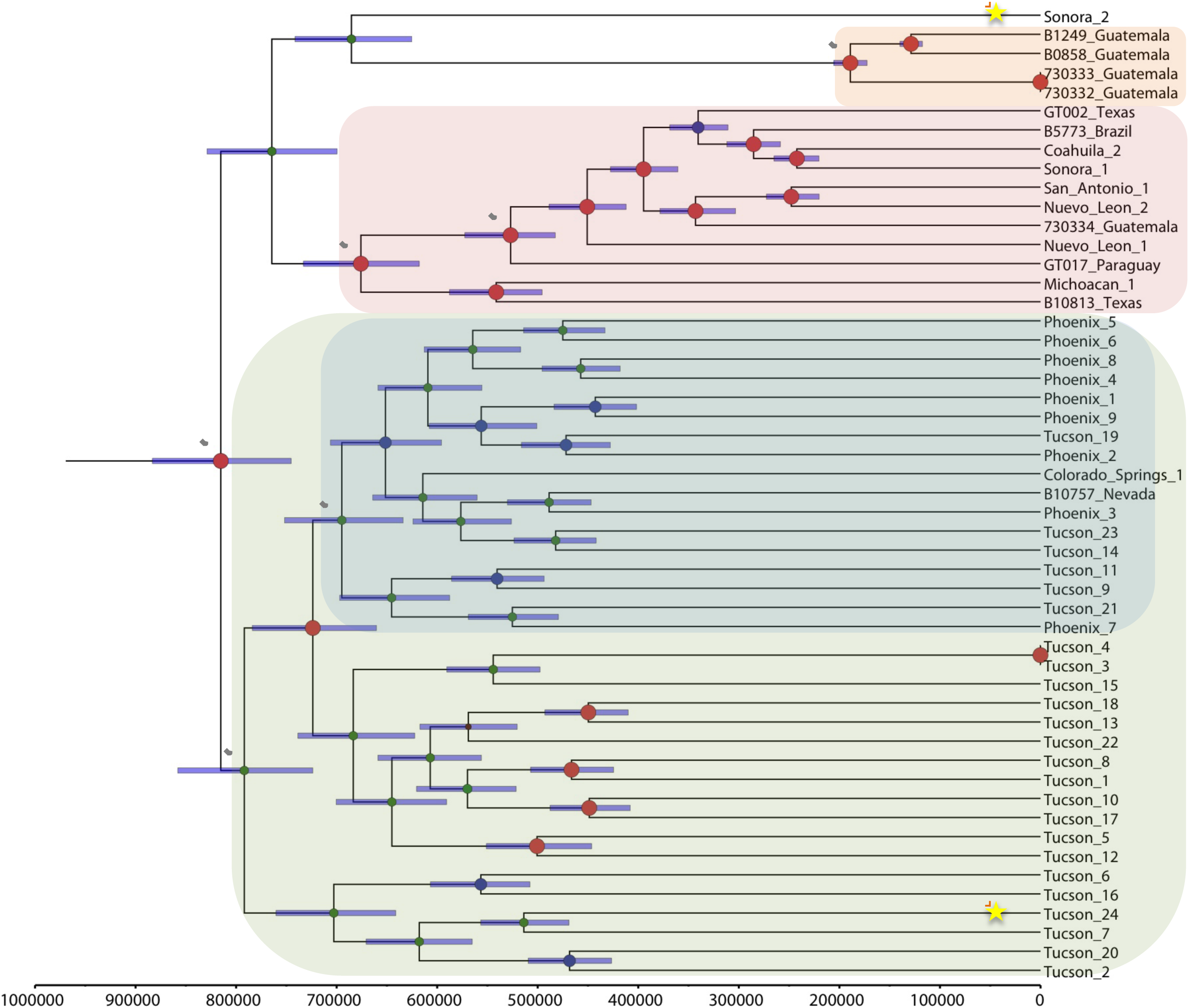
Bayesian phylogenetic analysis of *C. posadasii* isolates. BEAST 1.8.1 was used to produce a calibrated phylogeny, where the time to most recent common ancestor estimated for *C. posadasii* from the dual species analysis in Figure 1 was used for calibration here (TMRCA *C. posadasii:* 818,100 years ago). The analysis was performed on WGS SNP data from 51 C. *posadasii* genomes. Clades of interest are highlighted: green: Arizona, blue: Phoenix subclade of Arizona, orange: Guatemala, and pink: Texas-Mexico-South America. Stars highlight strains of interest while dotted circles represent TMRCA of interest. Posterior probabilities are indicated as node size. Purple node bars are shown for each node and are informative for the 95% confidence interval for the timing estimate. The timeline represent years before the present.

**Figure 3.**
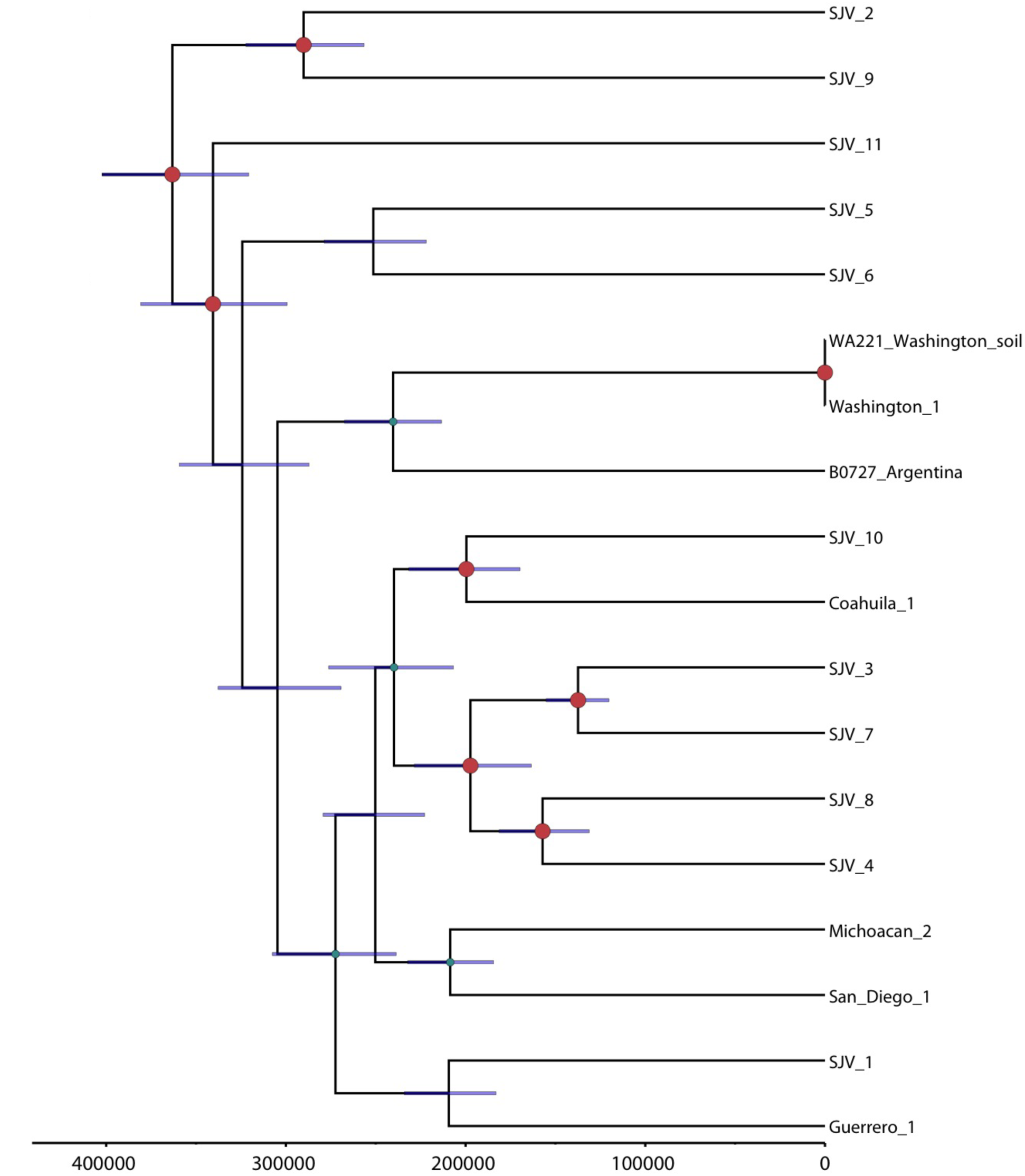
Bayesian phylogenetic analysis of *C. immitis* isolates. BEAST 1.8.1 was used to produce a calibrated phylogeny, where the time to most recent common ancestor estimated for *C. immitis* fi:Om the dual species analysis in Figure 1 was used for calibration here (TMRCA *C. posadasii:* 365,700 years ago). The analysis was performed onWG. SNP data from 18 C *posadasii* genomes. Posterior probabilities are indicated as node size. Purple node bars are shown for each node and are informative for the 95% confidence interval for the timing estimate. The timeline represent years before the present.

**Figure 4.**
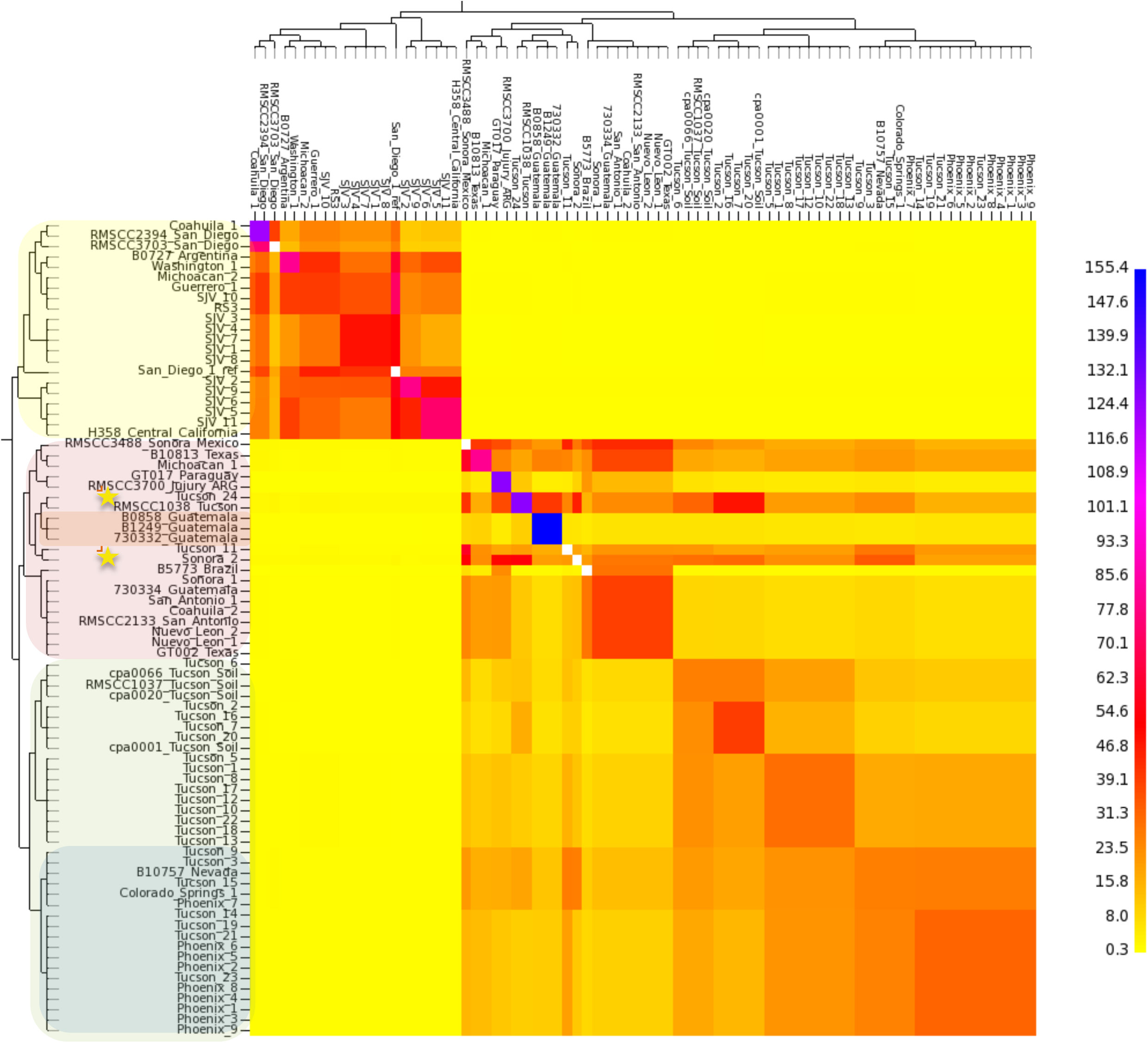
Genome sharing analysis of *Coccidioides* species complex. *fineStructure* analysis was performed using the SN. matrix developed for Supp. Figure 1. The SNPs from 66 *Coccidioides* spp. genomes were reduced to a pairwise similarity matrix, which was used to identify population structure based on shared haplotype regions of genome. The x-axis analysis represents the strain as a “recipient” and the y-axis represents the strain as a “donor” of genomic regions. The scale bar represents the number of shared genome regions with blue being the greatest amount of sharing and yellow being the least. The shading of isolates on the y-axis correlates with clades in Figure 1.

**Figure 5.**
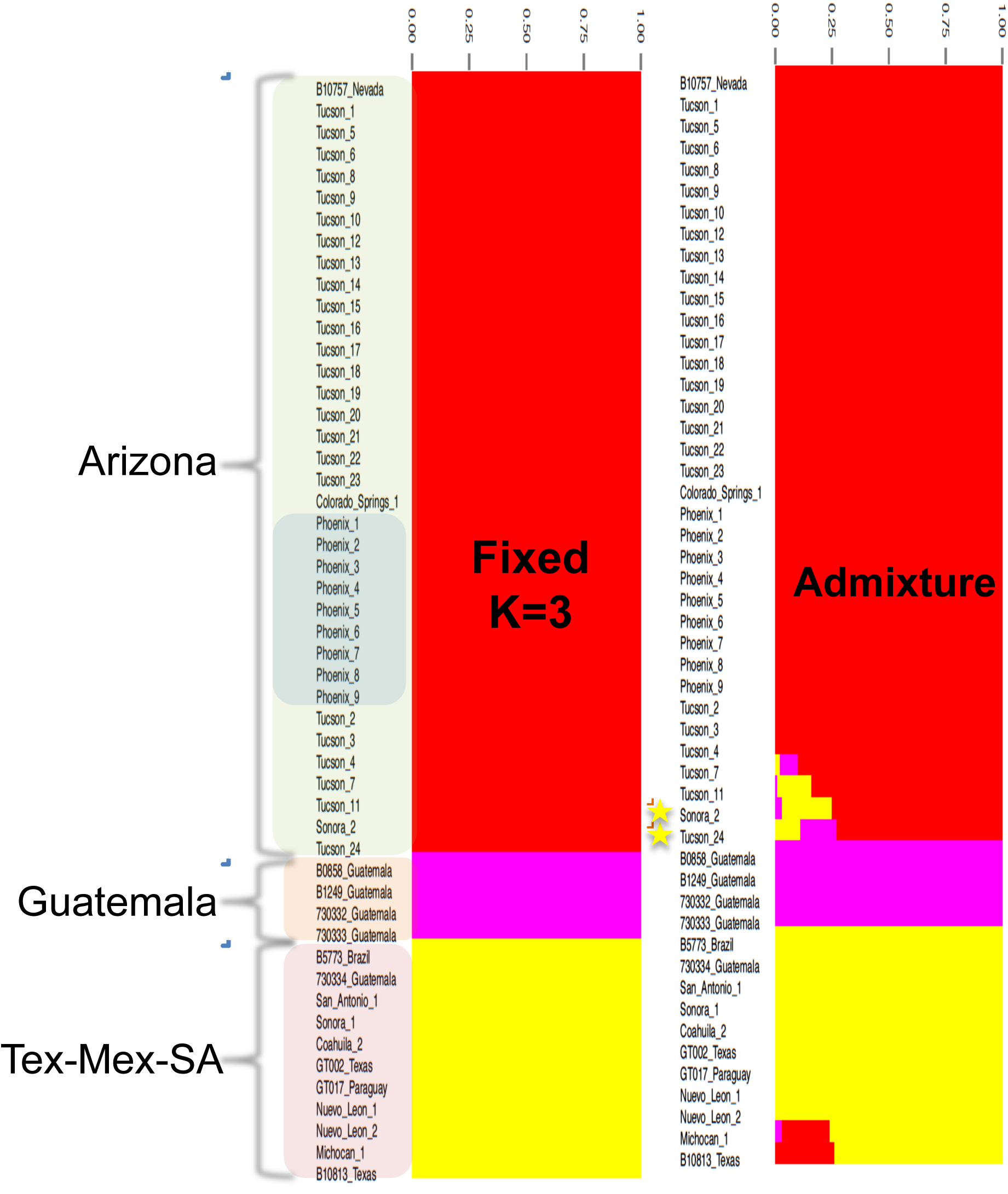
Population structure analysis of *C. posadasii*. Bayesian analysis of *C. posadasii* population structure was carried using BAPS 6.0 using 3 fixed genetically diverged groups previously established by phylogenetic inferences (**Supp. Figure 2**), using ten replicates. Admixture graphs of three identified *C. posadasii* mixtures (populations) was performed using 200 simulations and the percentage of genetic composition from each isolate was plotted. The shading of isolates in left column correlates with clades in Figure 1.

Within the Arizona population, several subclades were identified by the phylogenetic analyses and well supported by the ML bootstrap analysis (Figure S4). One subclade included all isolates from the Central Arizona (i.e., Phoenix) region as well as a small numbers of strains from Tucson, and the other two subclades were primarily comprised of the isolates from the Tucson region. However, the low Fst (0.043) and ΦPT (0.039) values and population genetics methods (Figures 4 and 5) suggest that these geographically distinct subpopulations are not genetically distinct.

The second *C. posadasii* population included almost all isolates from Texas, Mexico and South America, except for the “Sonora_2” strain, from northern Mexico, which appears as basal to this group in the maximum parsimony and neighbor network analysis (Supplemental Figures 3 and 5). The third population of *C. posadasii* included four Central American isolates from Guatemala that formed their own distinct population, supported by all analyses. A fifth Guatemalan isolate (i.e., “730334”) was isolated from a patient, who was infected while traveling in Texas, which is consistent with the placement of this isolate in the Texas/Mexico/S. America population in all phylogenetic analyses.

In addition, two strains, “Sonora_2” and “Tucson_24”, appear as distinct early lineages in the non-Arizona clades. Furthermore, when both *C. posadasii* and *C. immitis* isolates were included in the analyses, Bayesian as well as maximum likelihood methods identified “Sonora_2” and “Tucson_24” as possible basal lineages to the Guatemalan population (Figure 1 and Supp. Figure 4). However, when only *C. posadasii* strains were included in the Bayesian analysis, “Tucson_24” was placed within the Arizona population, while “Sonora_2” remained basal to the Guatemalan clade (Figure 2). Multiple phylogenetic methods using the same nucleotide substitution model produced phylogenies with slightly varying results, suggesting the observed incongruences are an artifact of differing algorithms implemented in differing phylogenetic methods.

### Coccidioides immitis

We identified 64,096 shared SNP loci (31,372 parsimony informative) among 18 analyzed *C. immitis* genomes. Bayesian (Figure 3), BAPS (Supplemental Figure 6), maximum likelihood analyses (Supplemental Figure 7), and maximum parsimony (Supplemental Figure 8) each displayed similar overall topologies for *C. immitis*, where generally each subpopulation or clade contained at least one member from the Central Valley of California (i.e., San Joaquin Valley). The Washington State (WA) and Argentina isolates typically grouped together. The Argentinian isolate came from a patient that was treated in Buenos Aries (not thought to be an endemic site and greater than 1200 km from the nearest known endemic zone for *C. posadasii*) and possibly represents an exposure outside of Argentina (although autochthonous South American exposures to *C. immitis* have been recently reported (20)). The Washington isolates were previously established to be recently endemic to the region (3) (and described as a distinct genetic subpopulation by BAPS analysis - Supplemental Figure 6). The strength of all of these analyses, however, is limited due to fewer *C. immitis* genomes in this study.

### Population genomics of and mating type distribution of Coccidioides

In order to explore previously identified genetic recombination (21,22) in the population analysis, we applied *fineStructure* and BAPS, model-based Bayesian approaches (23, 24) for incorporating recombination signal into population structure. The genus-level *fineStructure* analysis appropriately separated 81 *Coccidioides* genomes into the two species and assigned within-species “populations” or groups, related by shared genomic regions, which were similar although not identical to those described by other methods (Figure 4). Genomic isolation (shown as yellow on the heat map) was demonstrated for the two species; however, shared genetic history (showed by a gradual coloration on the heat map) was seen among members within each species. Several *C. posadasii* isolates displayed shared genomic space with members from outside their respective assigned groups, indicating the presence of historical admixture or incomplete (or recent) separation between groups within the species (Figures 4 and 5). Notably, evidence was provided for an extensive genome sharing history of both the “Tucson_24” and “Sonora_2” strains with the remainder of the *C. posadasii* isolates, providing additional evidence that these genomes are representative of older, more ancestral lineages, and explaining why different analyses group these strains differently (note: 1038_Tucson appears to be closely related to Tucson_24, however this is one of the previously Sanger-sequenced strains, and it is therefore not included in all analyses). The BAPS admixture analysis determined that admixture was present in these two strains, along with Tucson_11 and Tucson_7, as well as in B10813_Texas and Michocan_1 (Figure 5). Given the geographic separation, the identified admixture is likely indicative of lineage history rather than recent recombination between subpopulations. Both *fineStructure* and BAPS analyses highlighted the relative isolation of the Guatemalan Clade, as well as limited inter-population genome sharing of the individual strains from Brazil, Paraguay and Argentina, with the latter two displaying the most between-strain genomic sharing outside of the Guatemalan group.

Population structure of *C. immitis* reveals strong genetic isolation of WA isolates. In addition there is low genetic variation within the WA clade as represented by short internal branches. This characteristic is unique to the WA clade, and not shared within any other *C. immitis* or *C. posadasii* populations where each isolate appears to represent a single haplotype. Because recombination has been frequently reported for *Coccidioides*, we assessed the mating type distribution in the main clades of *Coccidioides*. The majority *C. immitis* and *C. posadasii* clades display similar ratios of the *MAT1-1* and the *MAT1-2* idiomorphs, suggesting random mating is possible (chi-square test, P values of > 0.05) (Supplementary Figure 10). However, we detected only a single idiomorph (*MAT1-2*) among six WA genomes sequenced so far, which is compatible with a clonal population. The uneven *MAT* idiomorph distribution was evidenced by a significant chi-square test (P=0.0143).

### Divergence and Most Recent Common Ancestor

Divergence analysis was calibrated using two previous estimates for the separation of the two species: 5.1 million years ago (MYA) (8) and 12.8 MYA (9). This analysis suggests that Arizona and non-Arizona *C. posadasii* subpopulations diverged between 820 thousand years ago (KYA) and 2.06 MYA, and the *C. immitis* populations diverged between 370 and 920 KYA, using the 5.1 MYA and 12.8 MYA calibration points, respectively. As the 5.1 MYA speciation estimate was derived from fossil data (8) rather than inferred from microsatellite mutation rate data, as was the case for the 12.8 MYA estimate (9), it was selected as the calibration point for estimating all other within-species divergence times (Table 2, Figure 2). The time to most recent common ancestor (TMRCA) of the Tex-Mex-SA population was estimated at 675 KYA (95% CI: 617-733 KYA), the first South American emergence is calculated at 527 KYA (95% CI: 483-573 KYA) and the Guatemalan MRCA emerged less than 190KYA (95% CI: 172-206 KYA). Using the combined runs for each species respectively and the 5.1 MYA calibration point, the *C. posadasii* estimated mutation rate was calculated to be 1.08*10^−9^ SNPs per base per year (95%CI: 9.9237*10^−10_-_^1.1752*10^−9^), and the estimated *C. immitis* mutation rate was 1.02*10^−9^ (95%CI: 9.0797*10^−10^-1.1431*10^−9^), similar to previous estimated mutation rates for fungi (11, 25).

**Table 2.**
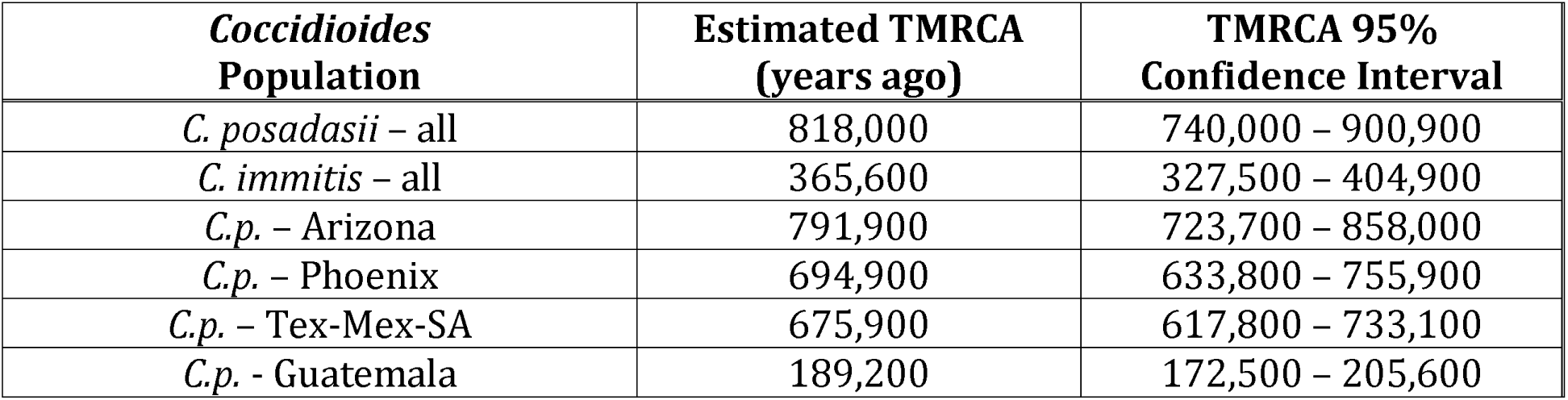
Estimated Time to Most Recent Common Ancestor for *Coccidioides* subpopulations, based on 5.1 MYA speciation calibration point.

| ***Coccidioides***<br>**Population** | **Estimated TMRCA**<br>**(years ago)** | **TMRCA 95%**<br>**Confidence Interval** |
| --- | --- | --- |
| *C. posadasii* – all | 818,000 | 740,000 – 900,900 |
| *C. immitis* – all | 365,600 | 327,500 – 404,900 |
| *C.p.* – Arizona | 791,900 | 723,700 – 858,000 |
| *C.p.* – Phoenix | 694,900 | 633,800 – 755,900 |
| *C.p.* – Tex-Mex-SA | 675,900 | 617,800 – 733,100 |
| *C.p.* - Guatemala | 189,200 | 172,500 – 205,600 |

## Discussion

Here we present results of the whole genome SNP analysis of *Coccidioides sp*. strains from geographically diverse locations that has further confirmed the presence of two species, *C. immitis* and *C. posadasii*, first established by Fisher *et al*. (2) and further described in subsequent studies (4,11). Using multiple methods of population and phylogenetic analyses, we provide evidence of further genetic subdivision within the major clades of *C. posadasii* and *C. immitis* and demonstrate the presence of genetically distinct subpopulations associated with specific geographic locales. Calibrated Bayesian analysis provides for a new understanding of the sequence of divergence events, particularly within *C. posadasii*. These data and analyses enhance our ability to conduct genomic epidemiology, make more informed hypotheses of likely ancestry and pathways of dispersal and better understand the changing distribution of *Coccidioides*.

The genetically differentiated Arizona and non-Arizona *C. posadasii* populations, first proposed by Fisher (9), are statistically distinct; and a third, previously unknown genetically isolated population in Guatemala, has now been identified. We are now able to identify multiple *C. posadasii* clades within Arizona and differentiate between isolates from the two major urban areas of Tucson (Southern AZ) and Phoenix (Central AZ), with the Phoenix isolates grouping in a single clade, although not forming a genetically isolated population (Figures 2, 4 and 5). Such fine scale resolution of subpopulations allows for a better understanding of originating source or location of non-endemic cases and recent emergences in new geographical regions. For example, epidemiological data strongly suggests that the “Guatemala_730334” case was not a Guatemalan exposure but rather an exposure in Texas, a supposition borne out by the genomic data. In the same respect “Colorado Springs_1” case was not likely a Colorado exposure, as there is no known endemic coccidioidomycosis occurring in that region of Colorado. However, whole genome SNP analyses place this strain within the Tucson/Phoenix subclade of the Arizona population of *C. posadasii*, likely representing the originating environmental source for that infection; unfortunately no travel history from this case was available for our study. In the same respect, the *C. immitis* cases from Lower Mexico (Guerrero and Michoacan) and Argentina (Buenos Aires), locales that are not known to support endemic populations of *C. immitis* (26,27), are possibly epidemiologically linked to their phylogeographic placement within one or more central California (e.g. San Joaquin Valley) clades. Such phylogeographic associations still require confirmation with local environmental sampling. For example, while the Washington isolates appear to fall within a larger San Joaquin Valley clade, it is now understood, through careful examination of both clinical and environmental isolates, that this clonal population emanates from a new endemic focus in southeastern Washington state (3,14), with possible original dispersal out of California’s Central Valley.

The resolution provided by whole genome SNP analysis also provides for an understanding of the possible ancestry of this fungus. Radiation of *C. posadasii* is estimated to have occurred ~450K years before the emergence of *C. immitis* (Figure 1, Table 2). This is clear evidence of an older and more diverse population, with *C. posadasii* likely being ancestral to *C. immitis*. Neafsey et al. (11) established that *C. posadasii* has a two-fold larger effective population size than *C. immitis*, suggesting that this was due to the larger geographic range, allowing for the development of more subpopulations. This, however, may also be explained by *C. posadasii* being an older population, with more time for mutation and divergence. It is of interest to consider that the center of diversity for *C. immitis* is found in the Central Valley of California, a geographic location borne of a large inland sea that began retreat during the Pleistocene (28) and most recently completely submerged by glacial runoff as long as 700,000 years ago (29). The estimated MRCA for *C. immitis* is 365K-920K years ago, depending on calibration model, which is subsequent to, or possibly coinciding with, the draining of the CA Central Valley. Isolation within glacial refugia has been proposed as a paleogeological factor impacting the historical continental dispersal of environmental fungi (30). The Sierra Nevada mountain range on the east side of the Central Valley, known to have been part of multiple glaciation events, could therefore have played a role as a geographic population barrier and a source for glacial refugia that subsequently drained in to the Central Valley, providing a possible paleogeographic cause for a speciation event.

Previous studies provided evidence of introgression between *C. posadasii* and *C. immitis*, which may have originated from a possible hybridization event in some Southern California and Baja Mexico isolates of *C. immitis* (11). In this study, we observed limited cross-species homoplasy, no identifiable genomic sharing (Figure 4), and an Fst value of >0.9; all of which are consistent with a limited population effect from hybridization. These data indicate that the two populations are largely reproductively isolated outside of the Southern Cahfornia region where suspected hybridization events are thought to have occurred (11). However, as the genomes from those previously identified introgressed strains were not of comparable quality to the genomes studied here, they were not analyzed in this study. Further evaluation of hybridization events and patterns of introgression is warranted.

The AZ clades of *C. posadasii* have an estimated MRCA approximately 115K years before the Tex-Mex-SA population MRCA (Figure 2), strongly suggesting that the Arizona group diverged earlier than non-Arizonan *C. posadasii* populations, although there is a slight overlap in confidence intervals (Table 2). A possible mechanism for the increased diversity within the AZ specific groups is increased recombination; however, only limited evidence of within haplotype sharing was seen with the *fineStucture* analysis (Figure 4), and each strain in this population corresponded to a single haplotype according to BAPS (Figure 5). When analyzed by itself, no admixture was identified within the AZ population (data not shown), further suggesting limited recombination is present among the genomes assessed. This limited genetic sharing may be a result of restricted recombination or historical mating events.

Both mating type loci were seen in roughly similar proportions in all geographic areas, except WA (Supplemental Figure 9), suggesting sexual recombination is possible throughout most of the endemic range. However, *Coccidioides* like other fungal genera may be impacted by a number of recombination/mating restrictions, both extrinsic (e.g. geographic separation) and intrinsic (e.g., lack of a need for regular sexual reproduction due to effective asexual mitosis strategy) (19). No laboratory-controlled genetic recombination with *Coccidioides* has been accomplished to date (31) and the role of, and barriers to, sexual reproduction in the natural setting remains unknown, similar to other environmental pathogenic fungi (30). The high genetic diversity observed within populations combined with observed homoplasy but limited detected recombination is more likely explained as incomplete lineage sorting. Ancestral polymorphisms likely remained through divergent events, accounting for varying alleles among related isolates but concurrent alleles in less closely related isolates (32). This would account for the long branch lengths, high homoplasy and little recombination seen among populations and individuals in the parsimony analyses.

Multiple clades occurring in Southern Arizona (i.e., Tucson), including one derived “younger” clade that contains all the Central Arizona (i.e., Phoenix) isolates, indicates that the Southern Arizona area is likely the source of the Central Arizona population, (Figure 2). In addition, maximum likelihood analysis indicates that the Tex-Mex-SA and the Guatemalan populations have also diverged from the Southern Arizona population (Supplemental figure 2); however other methods of phylogenetic reconstruction do not immediately support these relationships (Supplemental figures 1). A closer look at select strains may provide additional clues to the ancestral source of these populations. “Tucson_24” (along with the Sanger-sequenced “1038_Tucson” isolate) and “Sonora_2” (Northern Mexico) are placed within the non-Arizona clades in multiple Bayesian analyses (Figures 1,4,5) and may be ancestral to at least the Guatemalan population. “Sonora_2” and Texas isolate “B10813” appear to be the most basal members of the Tex-Mex-SA clade in the neighbor-net tree (Supplemental Figure 5), although greatly diverged from each other.

The same Texas isolate (“B10813”) and the “Michoacan_1” (Mexico) isolate are the two most basal to the Tex-Mex-SA clade in the Bayesian tree analyses (Figures 1 and 2) and in the maximum parsimony tree (where the Guatemala clade was used at the root) (Supplemental Figure 3). Additionally, “Tucson_24” and “Sonora_2” displayed the highest levels of admixture with all populations (Figure 5) and appeared to have the most “shared” haplotype space among all *C. posadasii* isolates in the *fineStructure* analysis of this clade (Figure 4), suggesting, perhaps, an ancestral genomic background for the species. The South American isolates appear to be derived in Tex-Mex-SA clade. The Argentina (“RMSCC_3700”) and Paraguay (“GT_1078”) isolates are closely related to each other and are closer to other Mexico strains than the Brazil strain (“B5773”), by both SNP distance and haplotype sharing, suggesting more than one independent founder events into South America.

The Guatemala clade is a genetically distinct subpopulation, and emerged much later than the Tex-Mex-SA population. While tropical Guatemala would typically be considered to be outside the arid endemic regions, multiple accounts of endemic transmission have been recorded in its arid Motagua River valley (27,33). These data lend evidence to a local distinct population of *Coccidioides* in Central America, with more recent divergence between individuals than seen in other locales. Evidence exists for an additional Central American population in the arid Comayagua Valley in Honduras (1,34), although no isolates were available for analysis in this study.

The phylogenetic clades or populations identified in the differing geographic regions likely reflect single or limited founder population events, followed by local evolution. Such events would conflict with the hypothesis of ongoing deposition of spores by wind as a likely mechanism for large-scale geographic dispersal (17), as continual dispersal would result in multiple distinct populations in each locale. For example, the grouping of central Arizona (Phoenix) isolates largely cluster in a single subclade of the southern Arizona population. The limited presence of Phoenix isolates outside this clade may represent instances of wind dispersal and/or exposure (i.e., infection in Tucson patient by wind-borne Phoenix-originating *Coccidioides* spores), although it is more plausible that patients infected in Phoenix are occasionally diagnosed in Tucson, and vice versa, as there is a high degree of travel between the two population centers. Cases of patients living in one endemic area but having exposure in another endemic area have been well documented (21,35).

A more likely hypothesis is that primary dispersal over large regions occurs through movement by mammalian hosts, similar to Fisher’s previously proposed mechanism for emergence of *C. posadasii* in South America during the Great American Biotic Interchange (GABI) via the Central American land bridge between the continents (9). The *Coccidioides* genomic adaptations to mammalian host (e.g., expansion of protease and keratinase gene families) (8, 36) and the hypothesis that the patchy distribution of *Coccidioides* in soil is due to the fungal association with mammal carcasses (i.e., dead hosts) and burrows (31,37) comports with a theory that distribution is related to distinct movement events of infected animals. Although multiple North American mammal species can be infected, rodents, canids and humans are highly susceptible to succumbing to the disease, allowing infecting strains to enter back into the soil upon death (1,38), and therefore may be considered the most likely “vectors” of *Coccidioides* from one locale to another. In South America, llamas and armadillos (39,40) are additionally considered highly susceptible and the latter being a known source for soil contamination and human exposure (41). The ecological importance of recent findings of infected bat populations in Brazil remains unknown (42).

The TMRCA analysis would suggest a separation of the *C. posadasii* subpopulations over the previous 800K years, the dispersal into South America no earlier than 575K years and the emergence in Guatemala less than 200K year ago. A closer look at the timing and composition of the known GABI events as compared to the timing of the *C. posadasii* southern dispersions is warranted. There are recorded limited mammal interchanges between the North and South American continents as early as 9 mya (43), however, animal exchanges did not begin in earnest until after the final settling of the Central American isthmus about 2.8 mya (44). Starting about 2.6 mya there were four significant exchanges of mammals (i.e., GABI 1-4) between North America and South America, typically separated by glacial retreat and other geo-climatic events (43). Humans are understood to have appeared in the Western Hemisphere and migrated to South America well after the last of the GABI events, likely less than 15,000 y.a. (45). The earliest estimated emergence of *C. posadasii* in South America (TMRCA = 527 kya) would have followed the GABI 3 event of multiple carnivores (including canid taxa) emigrating from North America to South America approximately 0.8M to 1.0 mya. The final GABI event (~125 kya) occurred subsequent to the establishment of the Guatemala *C. posadasii* population (TMRCA = ~189 kya). Of note, the earliest identification of the genus *Canis* in South America occurred at this time during GABI 4. The establishment of *C. posadasii* in Central America just prior to the GABI 4 event may then be related to the Central American “holding-pen” concept, where migrant mammalian taxa are thought to have established local populations and undergone provincial evolution prior to continuing their southward or northward expansions (43). Such an occurrence could account for a regional establishment of a distinct population of *C. posadasii* at this time. It is also possible that *Coccidioides* was more prevalent throughout Central America during the late Pleistocene epoch, where records show a vastly different climate and biotic landscape of very dry thorny-scrub and grasslands (46). The current known distribution of *C. posadasii* in Central America is in the dry Motagua River valley Guatemala and the dry regions of the Comayagua Valley of Honduras (33), perhaps the remaining vestiges of the historically dry *Coccidioides-supporting* landscape in the region. Lastly, the data presented here strongly suggest that humans were not the primary driver of *Coccidioides* dispersal into Central and South America, as had been previously postulated (9).

An updated *C. posadasii* dispersal model (Figure 6) would suggest that: A) the central Arizona clade originated from one of southern Arizona sub-populations, with all Arizona populations being 700-800K years old; B) Texas and Mexico populations also came from a founding population from the Southern Arizona-Sonora region, likely 675-700K years ago; C) Mexico (and possibly Texas) subsequently fed the South American populations (likely more than once), as early as 525K years ago; and D) the Guatemalan population independently, and more recently (<200K year ago), emerged from the Southern Arizona-Sonora region. The evidence of historical admixture of both “Tucson_24” and “Sonora_2” strains with the rest of the species, and the possible basal nature of the “Sonora_2” and/or “Tucson_24” strains to the Tex-Mex-SA and Guatemalan populations, suggests that these strains are from older lineages and could represent a historical southern Arizona-northern Mexico origin for the species.

**Figure 6.**
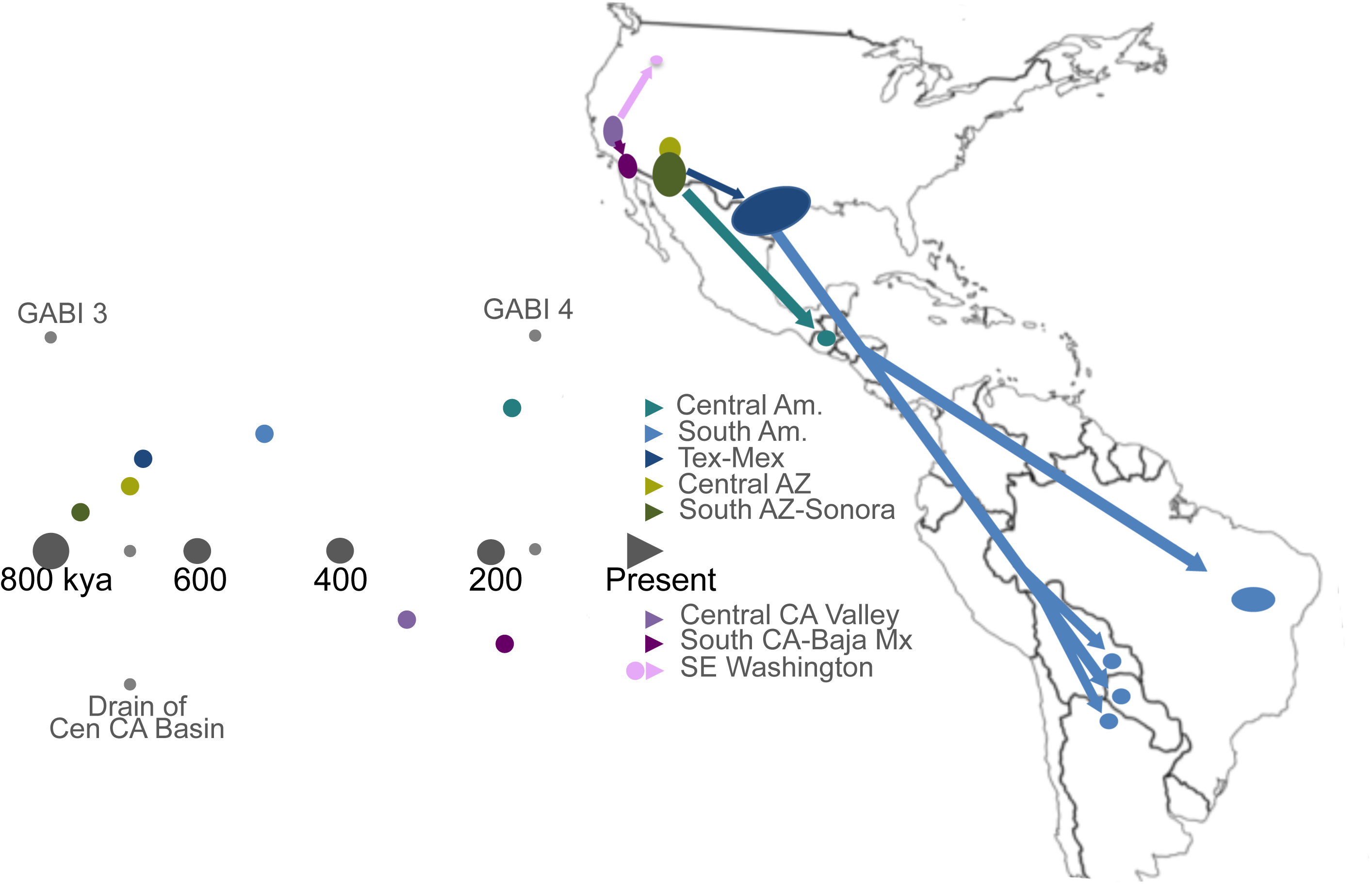
Model for *Coccidioides* dispersal in the Western Hemisphere. A proposed dispersal model for *C. posadasii* from a hypothetical founder population in Southern Arizona-Northern Mexico and *C. immitis* from a hypothetical founder population in California’s Central Valley is shown in conjunction with a timeline (labeled in thousands of years) that is color-coded to match dispersal paths and annotated with timing of Great American Biotic Interchange (GABI) events 3 and *4*, as well as the final Draining of the Central California Basin.

While no dispersal model for *C. immitis* is specifically proposed at this time, it is clear that much of the known diversity is found in the Central Valley of California (i.e. the San Joaquin Valley), with the WA clone being the likely most recent emigrant from this region. The Central Valley being historically submerged under inland sea and fresh water glacial lakes, and the final draining of this valley occurring an estimated 700K years ago, precedes the likely *C. immitis* species MRCA. Additional isolates from outside this region will need to be analyzed to better understand the genomic and dispersal history of this species.

We recognize the limitations of this study include sampling bias, especially with regards to analyzing fewer *C. immitis* and South American *C. posadasii* genomes, and hence a possibility for both over and under-representation of isolates from some geographic regions. Further conclusions regarding the *C. immitis* population structure should only be drawn from the inclusion of additional genomes from more locales. The vast majority of the isolates studied here originated from clinical samples. It is possible that environmental soil isolates may provide differing genotypes than those from infecting strains, although this was not seen in previous analyses (7), nor was this specifically seen with the inclusion of previously sequenced genomes from limited *C. posadasii* soil isolates (Supplemental Figure 1). Additionally, carefully collected environmental isolates would provide a more reliable representation of locally present strains (31). It is clear that future studies will require a more in depth analysis of environmental and clinical isolates. It is also important to note that early sequence data sets likely have less accuracy and coverage, affecting their utility for whole genome analyses, such as the ones conducted here. Current sequencing provides excellent depth and quality that should allow their use for studies well into the future. The use of numerous phylogenetic tools was employed to improve our understanding of the relationships between individuals and groups. It is understood that no single approach likely represents absolute truth, particularly for a dimorphic fungus with cryptic reproduction, however the combination of phylogenetic representations provided advances our understanding of potential historical relationships. Likewise, the use of Bayesian tools to determine mutation rates and times to most recent common ancestor does not likely represent actual truth, however, we employed a fossil-derived estimated calibration point for original speciation to provide a most likely representation of specific divergence points, rather than using an earlier calibration based on estimated microsatellite mutation rates. Additionally, we chose to infer molecular rates and divergence times from this calibration point rather than inferring mutation rates and coalescence from tip-dated sequences, which has been suggested to introduce positive-bias (47). As more genomes are sequenced from the regions of interest in this study, and additional calibration points are established (e.g., from ancient DNA analysis), the time to most recent common ancestor findings should continue to be re-assessed.

The findings of this study provide a strong argument for additional large-scale population level sequencing of *Coccidioides*, particularly for both clinical and soil isolates from under represented areas, especially Sonora/northern Mexico populations, to further understand the geographic extent of a possible southern Arizona-northern Mexico founding population. Even more importantly, our data demonstrate that local population structure does occur, even in recombining organisms, and that WGS approaches can be readily used for fungal molecular epidemiology of not only suspect clonal outbreaks, but also for linking cases to likely exposure sites and better understanding the patterns of emergence and dispersal.

## Methods

### Isolates Selected for Sequencing

Isolates of 50 C. *posadasii* and 18 C. *immitis* were selected from multiple repositories among the collaborators on this study, to include primarily geographic diversity (Table S1) and extensive samples for regions of high disease burden (i.e., Southern Arizona). Isolate locales are based on available metadata and may represent location of sample isolation, laboratory that stored isolate, or home of patient.

### Genome Sequencing

The genomes of 18 *C. immitis* and 50 *C. posadasii* isolates were sequenced using Illumina HiSeq and MiSeq sequencing platforms, as previously described (14,48). High molecular weight DNA was extracted using the ZR Fungal/Bacterial DNA Mini Prep kit (Zymo Research, Irvine, CA, USA, catalog #D6005). DNA samples were prepared for paired-end sequencing using the Kapa Biosystems library preparation kit (Woburn, MA, USA, catalog #KK8201) protocol with an 8bp index modification. Briefly, 2 μg of dsDNA sheared to an average size of 600 bp was use for the Kapa Illumina paired-end library preparation, as described by the manufacturer. Modified oligonucleotides (Integrated DNA Technologies, Coralville, ΙΑ, USA) with 8bp indexing capability (49), were substituted at the appropriate step. Prior to sequencing, the libraries were quantified with qPCR on the 7900HT (Life Technologies Corporation, Carlsbad, CA, USA) using the Kapa Library Quantification Kit (Woburn, MA, USA, catalog #KK4835). Libraries were sequenced to a read length of 100bp or 150bp on the Illumina HiSeq or to 250bp on the Illumina MiSeq. All WGS data files have been deposited in the NCBI Sequence Read Archive (http://www.ncbi.nlm.nih.gov/bioproject, project # PRJNA274372)

### Genome Assembly

The “San Diego_1” (*C. immitis)* and “B10813_Tx” (*C. posadsasii)* sequenced genomes were both *de novo* assembled using the SPAdes assembler v2.5.0 (50). The “San Diego_1” assembly was used as the reference for the “All Species” and *C. immitis* SNP matrices and the “B10813_Tx” assembly was used as the *C. posadasii* SNP matrix (see SNP Variant Detection below)

### SNP Variant Detection

Illumina read data were aligned against the respective reference assemblies using Novoalign 3.00.03 (www.novocraft.com) and then SNP variants were identified using the GATK Unified Genotyper v2.4 (51). SNP calls were then filtered using a publically available SNP analysis pipeline (http://tgennorth.github.io/NASP/), as previously described (48), to remove positions with less than 10x coverage, with less than 90% variant allele calls, or identified by Nucmer as being within duplicated regions in the reference. SNP matrices were produced for the “all species”, *C. posadasii*, and *C. immitis* analyses.

### Phylogenetic Analysis

To understand relationships between isolates, we conducted multiple phylogenetic analyses including parsimony, likelihood and Bayesian inference using whole genome SNP matrices from a total of 86 genomes (60 *C. posadasii* and 26 *C. immitis*), which included newly sequenced and previously published genomes (Table S1). Maximum parsimony SNP trees, based on each of the SNP matrices, were constructed, using PAUP*v.4.0b10 (52) and visualized in Figtree v.1.3.1 (http://tree.bio.ed.ac.uk/software/figtree/). The maximum parsimony all species *Coccidioides* tree was mid-point rooted and used “San Diego_1” as a reference. The maximum parsimony *C. immitis* tree was rooted using “Coahuila_1”, based on its basal position in the all species tree, and “San Diego_1” was used as the alignment reference. The maximum parsimony *C. posadasii* tree was rooted using the Guatemala clade, again based on its basal position in the all species tree, and “B10813_Tx” was used as the alignment reference. Due to likely recombination present in the SNP data, a neighbor joining splits tree network or “neighbor-net” tree (53) was drawn to visualize genome sharing between isolates in each of the species trees. The neighbor-net was drawn using SplitsTree4 (54) with the uncorrected P distance transformation, as previously described (48). Maximum likelihood trees were produced with a rapid and effective stochastic algorithm implemented in IQ-Tree (55). The inferred nucleotide substitution model was TIM3+ASC+R10 for *C. posadasii* and the TVM+ASC+R7 for *C. immitis* and 1,000 non-parametric bootstrap pseudoreplicates were performed for branch confidence (56). BEAST v1.8.0 (57) and BAPS (24) were used to infer population structure and produce Bayesian phylogenetic trees; and BEAST was additionally used for mutation rate and time to most recent common ancestor analysis (TMRCA) (see a description of BEAST analyses below). Due to incongruences observed when comparing the Bayesian, maximum likelihood and maximum parsimony trees, we separately compared the same models that were used in the BEAST Bayesian analysis and the IQ-Tree maximum likelihood analysis in order to test the affect of model selection on the phylogenetic topologies. Maximum likelihood trees produced using both the HKY+G and the GTR+G models were identical and both demonstrated the same phylogenetic incongruences when compared to the BEAST trees (data not shown). The observed phylogenetic differences are therefore more likely a product of differing methods rather than nucleotide substitution model selection.

For the BAPS analysis, population assignments were assessed using the ‘Fixed-K Model’. The number of populations (genetic mixture) were tested from KΦ=Φ2 to K=15 for *C. posadasii* and K=2 to K=5 for *C. immitis* using 10 replicates for each tested K. Coccidioides admixed ancestry from previously inferred partitions (K=3 for both species) were conducted using 200 iterations. Admixture results were bar-plotted with help of STRUCTURE PLOT (58). Additionally, we tested the Arizona population of *C. posadasii* individually to assess admixture occurring within this population using BAPS. No evidence of admixture or recombination within this population was identified (data not shown). *fineStructure* population analysis (23) was implemented on the all species SNP matrix in order to infer recombination and population structure based on the presence or absence of shared genomic haplotype regions, as described elsewhere (48). In brief, the SNP matrix was reduced to a pairwise similarity matrix using Chromopainter, within the *fineStructure* program, employing the linkage model, where the underlying model assumes individuals within populations will share more regions of their genome with each other and have a similar amount of admixture with individuals from different populations. Identified populations are merged one at a time, at each step using the most likely population merge, to relate populations to each other through a tree.

In order to determine the amount of between-population and within-population variance we applied an “analysis of molecular variance” (AMOVA) using the GenAlEx program (59). The AMOVA produces an Fst score and a ΦPT score, which is an analogue of Wright’s Fst. A ΦPTϏ=Ϗ0 is considered indicative of no genetic difference among populations and ΦPTϏ=Ϗ1 indicates 100% genetic variance. MEGA (60) was used to calculate the average SNP distance within each identified population, (i.e., the number of base substitutions per site from averaging over all genome pairs within each population).

### Divergence Time Analysis

To estimate evolutionary rates for all of *Coccidioides*, as well as divergence times for *immitis* and *posadasii*, we employed a Bayesian molecular clock method as implemented in the BEAST v1.8.0 software package (57). Model selection analyses were carried out in PAUP* version 4.0a142 for the complete *Coccidioides* dataset and for each of the two species separately, where the Bayesian Information Criterion results were used to determine the best fitting models. While the model selection results were in concordance with the maximum likelihood model selection, BEAST currently has a limited selection of nucleotide substitution models available. Of the models available to set in BEAST, the GTR+G model was found to be best fitting for both the *Coccidioides* and *C. immitis* datasets, while the HKY+G model was determined to be best fitting for the *C. posadasii* dataset. Because only variable sites were included in this analysis, we corrected for the invariant sites by specifying a Constant Patterns model in the Patterns List of the BEAST xml file. For each dataset, the Constant Patterns model includes the number of constant A’s, C’s, T’s, and G’s. Given the high number of SNPs that compose the dataset, path sampling (61) and stepping stone (62) sampling marginal likelihood estimators were computationally prohibitive, so we instead employed the Akaike’s Information Criterion (AICM) to determine the best fitting clock and demographic model combinations (63,64). Model comparison analyses indicated that the combination of the strict molecular clock and the constant population models best fit all three datasets.

The strict clock model was used to infer the timescale and mutation rates through the incorporation of two separate internal calibrations on the node defining the speciation event. The first speciation calibration date, from Sharpton et al. (8), was estimated to be 5.1MYA through the use of a prior fossil-based calibration point. Previously, Fisher et al. (9) calculated a deeper speciation time of 12.8 MYA by determining microsatellite flanking sequence genetic distances. A normal prior age distribution with a standard deviation (sd)

0.25MYA was used for each of the full *Coccidioides* analyses. For the within species BEAST analyses, the mean TMRCA timepoints from the 5.1MYA run for *C. immitis* (normal prior age distribution, mean: 365,700 years ago, sd: 30,000) and *C. posadasii* (normal prior age distribution, mean: 818,100 years ago, sd: 50,000) were used as a calibration points, respectively. Importantly, the normal prior age distribution was applied to all calibration points in this study, as calibrations based on molecular studies or those that are secondary calibrations have added uncertainty, and that uncertainty in age is equally distributed about the mean.

Visual trace inspection and calculation of effective sample sizes was conducted using Tracer (65), confirming MCMC mixing within chains, and also among chains for the full *Coccidioides* runs. However, in agreement with the low consistency indices calculated during the parsimony analyses, we did not visualize among chain convergence within species. Consistent with high levels of recombination, the lack of among chain convergence indicates that multiple phylogenies can explain the evolutionary history within *Coccidioides* species. For each dataset, four independent Markov chain Monte Carlo (MCMC) chains were run for 100 million generations each, with parameters and trees drawn from the posterior every 10,000^th^ step. LogCombiner (57) was used to merge the samples from each chain. For each of the full *Coccidioides* analyses and for the *C. immitis* analysis, the first 20% of each chain was discarded as burn-in, and then each chain was resampled every 30,000^th^ step.

For the *C. posadasii* analysis, within chain convergence occurred later in two of the four runs, and the first 50% and 80% of the runs were discarded, and all four chains were resampled every 20,000^th^ step. The posterior mean and 95% confidence intervals have been reported for evolutionary rates and time to most recent common ancestor estimates for select nodes.

### Mating Type locus distribution

The *MAT1-1* gene from *C. immitis* (EF472259.1) and *MAT1-2* gene from *C. posadasii* (EF472258.1) were used as query sequences for sexual idiomorphic identification in *Coccidioides* (66). Protein sequences from *MAT* genes were searched against each assembled genome herein analyzed via tBLASTn tool (67) and retrieved alignments were manually inspected in order to check the completeness of each *Coccidioides MAT* gene identified. The idiomorphic distributions were assigned to each clade deduced by ML tree. Mating type distribution significances were tested for deviation from the expected *MAT* ratio of 1:1 using a chi-square test using Microsoft Excel. Uneven *MAT* distribution were considered significant those with P < 0.05.

## Acknowledgements

We thank Blanca Samayoa, Dalia Lau, Ligia Figueroa, Danicela Mercado and Brenda Guzman for assistance with sample collection and shipment. We thank the Monsoon High Performance Computing Cluster Program, at Northern Arizona University for support with BEAST analyses.

The authors declare no conflicts of interest.

## Author contributions

D.M.E., C.C.R., E.M.D., J.M.S., B.B., and P.K. designed research; D.M.E., C.C.R., C.M.H., M.T., E.M.D., A.P.L., L. G., E.A., H.L., V.W., K.K., G.R.T., T. C., B.B. and P.K. performed research; D.M.E., C.C.R., C.M.H., M.T., A.P.L., B.B., and P.K. analyzed data; D.M.E., C.C.R., C.M.H., P.K. and A.P.L.wrote the paper

## Supplemental Figures

**Supp. Figure 1. Phylogenetic analysis of *C. immitis* and *C. posadasii* isolates from all known endemic regions**. Maximum parsimony phylogenetic analysis was performed on WGS SNP data from 81 *Coccidioides* genomes including 12 publically available genomes: 22 *C. immitis* and 59 *C. posadasii*. The shading of clades correlates with clades in Figure 1. The analysis identified 128,871 shared SNPs, with 70,419 being parsimony informative, with a consistency index (CI) of 0.358 and a retention index (RI) of 0.847. The tree shown is midpoint rooted. Branch lengths represent numbers of SNPs between taxa, with the unit bar in the figure.

**Supp. Figure 2. Maximum likelihood SNP phylogenetic analysis of *C. immitis* and *C. posadasii* isolates**. Maximum likelihood analysis was performed on WGS SNP data from 69 *Coccidioides* genomes using the TIM3 + ASC + R10 model with the program IQ-TREE with 1000 boostrap support. The shading of clades correlates with clades in Figure 1. The tree is mid-point rooted and San Diego_1 assembly was used as the reference.

**Supp. Figure 3. Maximum Parsimony phylogenetic analysis of *C. posadasii* with publically available genomes**. Maximum parsimony phylogenetic analysis was performed on WGS SNP data from 58 *C. posadasii* genomes, including 7 publically available genomes. The shading of clades correlates with clades in Figure 1. The analysis identified 142,261 total SNPs, with 67,166 parsimony informative SNPs, and a consistency index (CI) of 0.228 and a retention index (RI) of 0.36. The tree shown is rooted using the Guatemalan clade based on Supp. Figure 1. The unit bar in the figure represents SNPs.

**Supp. Figure 4. Maximum Likelihood phylogenetic analysis of *C. posadasii* isolates**. Maximum likelihood analysis was performed on WGS SNP data from 51 C. *posadasii* genomes. IQ TREE identified the TVM + ASC + R9 model as the correct model to use and 1000 bootstrap support was performed. The shading of clades correlates with clades in Figure 1. Numbers on branches represent bootstrap values. The tree is midpoint rooted.

**Supp. Figure 5. Phylogenetic network of *C. posadasii***. *A* neighbor-net representation of the relationships among the 51 C. *posadasii* isolates in Figure 3 based on SNP data, using the uncorrected P distance transformation. Each band of parallel edges indicates a split. Splits of major phylogeographically clustered subpopulations are shaded according to the key.

**Supp. Figure 6. Population structure analysis of *C. immitis* isolates**. Bayesian analysis of *C. immitis* population structure was carried using BAPS 6.0 using 2 fixed genetically diverged groups previously established by phylogenetic inferences (**Supp. Figure 2**), using ten replicates. Admixture graphs of the two identified *C. immitis* mixtures (populations) was performed using 200 simulations and the percentage of genetic composition from each isolate was plotted.

**Supp. Figure 7. Maximum Likelihood phylogenetic analysis of *C. immitis* isolates**. Maximum likelihood analysis was performed on WGS SNP data from 18 *C. immitis* isolates using the TVM + ASC + R7 model with 1000 bootstrap support. Numbers on branches represent bootstrap values. The tree was midpoint rooted.

**Supp. Figure 8. Maximum Parsimony phylogenetic analysis of *C. immitis* isolates**. Maximum parsimony analysis was performed on WGS SNP data from 18 *C. immitis* isolates. A total of 64,096 SNPs were identified, 31,372 were parsimony informative. The CI value was 0.346 and the RI was 0.384. The tree was rooted using Coahuila 1, based on the tree from Supp. Figure 1.

**Supp. Figure 9. Mating-type distribution of *Coccidioides* subpopulations**. We analyzed the mating type background from each available *Coccidioides* genomes sequenced so far. The *MAT1-1* gene from *C. immitis* (EF472259.1) and *MAT1-2* gene from *C. posadasii* (EF472258.1) were used as query sequences for sexual idiomorphic identification in *Coccidioides*. Mating types were counted for each *Coccidioides* diagnosed clades diagnosed by plylogenomic analysis. Mating type distribution significances were tested for deviation from the expected *MAT* ratio of 1:1 using a chi-square test. Uneven *MAT* distribution were considered significant those with P < 0.05.

## References

1. Brown J, Benedict K, Park BJ, Thompson GR 3rd. 2013. Coccidioidomycosis: epidemiology. Clin Epidemiol 5:185–97.

2. Fisher MC, Koenig GL, White TJ, Taylor JW. 2002. Molecular and phenotypic description of *Coccidioides posadasii* sp nov., previously recognized as the non-California population of *Coccidioides immitis*. Mycologia 94:73–84.

3. Marsden-Haug N, Hill H, Litvintseva AP, Engelthaler DM, Driebe EM, Roe CC, Ralston R, Hurst S, Goldoft M, Gade L, Wohrle R, Thompson GR, Brandt ME, Chiller T. 2014. *Coccidioides immitis* identified in soil outside of its known range — Washington, 2013. MMWR 63:450–450

4. Barker BM, Jewell KA, Kroken S, Orbach MJ. 2007. The population biology of *Coccidioides:* epidemiologic implications for disease outbreaks. Ann N Y Acad Sci 1111:147–63.

5. Cox RA, Magee DM. 2004. Coccidioidomycosis: host response and vaccine development. Clin Microbiol Rev 17:804–39.

6. Pappagianis, D. 1967. Epidemiological aspects of respiratory mycotic infections. Bacteriol Rev. 31:25–34.

7. Barker BM, Tabor JA, Shubitz LF, Perrill R, Orbach MJ. 2012. Detection and phylogenetic analysis of *Coccidioides posadasii* in Arizona soil samples. Fungal Ecol 5:163–176

8. Sharpton TJ, Stajich JE, Rounsley SD, Gardner MJ, Wortman JR, Jordar VS, Maiti R, Kodira CD, Neafsey DE, Zeng Q, Hung CY, McMahan C, Muszewska A, Grynberg M, Mandel MA, Kellner EM, Barker BM, Galgiani JN, Orbach MJ, Kirkland TN, Cole GT, Henn MR, Birren BW, Taylor JW. 2009. Comparative genomic analyses of the human fungal pathogens *Coccidioides* and their relatives. Genome Res 19:1722–31.

9. Fisher MC, Koenig GL, White TJ, San-Blas G, Negroni R, Alvarez IG, Wanke B, Taylor JW. 2001. Biogeographic range expansion into South America by *Coccidioides immitis* mirrors New World patterns of human migration. Proc Natl Acad Sci US. 98:4558–4562.

10. Engelthaler DM, Balajee SA. Forensics and Epidemiology of Fungal Pathogens, p 297–313. *In* Budowle, Schutzer, Breeze, Keim, and Morse (ed). Microbial Forensics, 2010. Elsevier Press, San Diego, CA.

11. Neafsey DE, Barker BM, Sharpton TJ, Stajich JE, Park DJ, Whiston E, Hung CY, McMahan C, White J, Sykes S, Heiman D, Young S, Zeng Q, Abouelleil A, Aftuck L, Bessette D, Brown A, FitzGerald M, Lui A, Macdonald JP, Priest M, Orbach MJ, Galgiani JN, Kirkland TN, Cole GT, Birren BW, Henn MR, Taylor JW, Rounsley SD. 2010. Population genomic sequencing of *Coccidioides* fungi reveals recent hybridization and transposon control. Genome Res 20:938–46.

12. Engelthaler DM, Chiller T, Schupp JA, Colvin J, Beckstrom-Sternberg SM, Driebe EM, Moses T, Tembe W, Sinari S, Beckstrom-Sternberg JS, Christoforides A, Pearson JV, Carpten J, Keim P, Peterson A, Terashita D, Balajee SA. 2011. Next-generation sequencing of *Coccidioides immitis* isolated during cluster investigation. Emerg Infect Dis 17:227–232.

13. Etienne KA, Gillece J, Hilsabeck R, Schupp JM, Colman R, Lockhart SR, Gade L, Thompson EH, Sutton DA, Neblett-Fanfair R Park BJ, Turabelidze G, Keim P, Brandt ME, Deak E, Engelthaler DM. 2012. Whole genome sequence typing to investigate the *Apophysomyces* outbreak following a tornado in Joplin, Missouri, 2011. PLoS One 7:e49989.

14. Litvintseva AP, Marsden-Haug N, Hurst S, Hill H, Gade L, Driebe EM, Ralston C, Roe C, Barker BB, Goldoft M, Keim P, Wohrle R, Thompson GR III, Engelthaler DM, Brandt ME, Chiller T. 2015. Valley Fever: Finding new places for an old disease: *Coccidioides immitis* found in Washington State soil associated with recent human infection. Clin Infect Dis 60:e1–3

15. Litvintseva AP, Hurst S, Gade L, Frace MA, Hilsabeck R, Schupp JM, Gillece JD, Roe C, Smith D, Keim P, Lockhart SR, Changayil S, Weil MR, MacCannell DR, Brandt ME, Engelthaler DM. 2014. Whole genome analysis of *Exserohilum rostratum* from the outbreak of fungal meningitis and other infections. J Clin Microbiol 52:3216–22

16. Fisher FS, Bultman MW, Johnson SM, Pappagianis D, Zaborsky E. 2007. *Coccidioides* niches and habitat parameters in the southwestern United States: a matter of scale. Ann NY Acad Sci. 1111:47–72.

17. Pappagianis D, Einstein H. 1978. Tempest from Tehachapi takes toll or *Coccidioides* conveyed aloft and afar. West J Med 129:527–530.

18. Jewell K, Cheshire R, Cage GD. 2008. Genetic diversity among clinical *Coccidioides* spp. isolates in Arizona. Med Mycol 46:449–55.

19. Taylor JW, Hann-Soden C, Branco S, Sylvain I, EUison CE. 2015. Clonal reproduction in fungi. Proc Natl Acad Sci USA. 112:8901–8

20. Canteros CE, Vélez H A, Toranzo AI, Suárez-Alvarez R, Tobón O Á, Del Pilar Jimenez A M, Restrepo M Á. 2015. Molecular identification of *Coccidioides immitis* in formalin-fixed, paraffin-embedded (FFPE) tissues from a Colombian patient. Med Mycol 53: 520–7.

21. Taylor JW, Geiser DM, Burt A, Koufopanou V. 1999. The evolutionary biology and population genetics underlying fungal strain typing. Clin Microbiol Rev 12:126–46.

22. Fisher MC, Rannala B, Chaturvedi V, Taylor JW. 2002. Disease surveillance in recombining pathogens: Multilocus genotypes identify sources of human *Coccidioides* infections. Proc Natl Acad Sci US. 99:9067–9071

23. Lawson DJ, Hellenthal G, Myers S, Falush D. 2012. Inference of population structure using dense haplotype data. PLoS Genet 8:e1002453.

24. Corander J, Marttinen P, Sirén J, Tang J. 2008. Enhanced Bayesian modelling in BAPS software for learning genetic structures of populations. BMC Bioinform 9:539.

25. Lynch M. 2010. Evolution of the mutation rate. Trends Genet 26:345–352.

26. Castañón-Olivares LR, Güereña-Elizalde D, González-Martínez MR, Licea-Navarro AF, González-González GM, Aroch-Calderón A. 2007. Molecular identification of *Coccidioides* isolates from Mexican patients. Ann N Y Acad Sci 1111:326–35.

27. Sifuentes-Osornio J, Corzo-León DE. Ponce-de-León LA. 2012. Epidemiology of Invasive Fungal Infections in Latin America. Curr Fungal Infect Rep 6:23–34.

28. Page RW. 1986. Geology of the fi-esh ground-water basin of the Central Valley, California, with texture maps and sections. USGS Prof Paper 1401:C1–C54.

29. Sarna-Wojcicki AM, Meyer CE, Bowman HR, Hall NT, Russell PC, Woodward MJ, Slate JL. 1985. Correlation of the Rockland ash bed, a 400,000-year-old stratigraphic marker in northern California and western Nevada and implications for middle Pleistocene paleogeography of central California. Quatern Res 23:236–57.

30. Gladieux P, Wilson BA, Perraudeau F, Montoya LA, Kowbel D, Hann-Soden C, Fischer M, Sylvain I, Jacobson DJ, Taylor JW. 2015. Genomic sequencing reveals historical. demographic and selective factors associated with the diversification of the fire-associated fungus Neurospora discreta. Mol Ecol 24:5657–5675.

31. Lewis ER, Bowers JR, Barker BM. 2015. Dust devil: the life and times of the fungus that causes valley Fever. PLoS Pathog 11: e1004762.

32. Stewart JE, Timmer LW, Lawrence CB, Pryor BM, Peever TL. 2014. Discord between morphological and phylogenetic species boundaries: incomplete lineage sorting and recombination results in fuzzy species boundaries in an asexual fungal pathogen. BMC Evol Biol 14:38.

33. Mayorga, RP and Espinoza H. 1970. Coccidioidomycosis in Mexico and Central America. Mycopathologia et Mycologia apphcata 40:13–23.

34. Mayorga, RP. 1967. Coccidioidomycosis in Central America, p 287–291 *In* Ajello L (ed). Coccidioidomycosis. The University of Arizona Press. Tucson, Arizona.

35. Burt AD, Carter A, Koenig GL, White TJ, Taylor JW. 1996. Molecular markers reveal cryptic sex in the human pathogen *Coccidioides immitis*. Proc Natl Acad Sci US. 93:770–773.

36. Whiston E, Taylor JW. 2015. Comparative Phylogenomics of Pathogenic and Non-pathogenic Species. G3 pii: g3.115.022806.

37. Whiston E, Taylor JW. 2014. Genomics in *Coccidioides:* insights into evolution, ecology, and pathogenesis. Med Mycol 52:149–55.

38. Sorensen RH. 1964. Survival characteristics of mycelia and spherules of coccidioides immitis in a simulated natural environment. Am J Hyg 80:275–85.

39. Diab S, Johnson SM, Garcia J, Carlson EL, Pappagianis D, Smith J, Uzal FA. 2013. Case report: Abortion and disseminated infection by *Coccidioides posadasii* in an alpaca *{Vicugna pacos*) fetus in Southern California. Med Mycol Case Rep 2:159–62.

40. Eulálio KD, de Macedo RL, Cavalcanti MA, Martins LM, Lazéra MS. Wanke B. 2001. *Coccidioides immitis* isolated from armadillos *[Dasypus novemcinctus)* in the state of Piauí, northeast Brazil. Mycopathologia 149:57–61.

41. Brillhante RS, Moreira Filho RE, Rocha MF, Castelo-Branco Dde S, Fechine MA, Lima RA, Picanço YV. Cordeiro Rde A, Camargo ZP, Queiroz JA, Araujo RW, Mesquita JR, Sidrim JJ. 2012. Coccidioidomycosis in armadillo hunters from the state of Ceará, Brazil. Mem Inst Oswaldo Cruz. 107:813–5.

42. Cordeiro RA, e Silva KR, Brilhante RS, Moura FB, Duarte NF, Marques FJ, Cordeiro Rde A, Filho RE, de Araújo RW. Bandeira Tde J, Rocha MF, Sidrim JJ. 2012. *Coccidioides posadasii* infection in bats, Brazil. Emerg Infect Dis 18: 668.

43. Woodborne MO. 2010. The Great American Biotic Interchange: Dispersals, Tectonics, Climate, Sea Level and Holding Pens. J Mamm Evol 17:245–264.

44. Bartoli G, Sarnthein M, Weinelt M, Erlenkeuser H, Garbe-Schönberg D, Lea DW. 2005. Final closure of Panama and the onset of northern hemisphere glaciation. Earth Planet Sci Lett 237:33–44.

45. Waters MR, Stafford TW Jr, Kooyman B, Hills LV. Late Pleistocene horse and camel hunting at the southern margin of the ice-free corridor: reassessing the age of Wally's Beach, Canada. Proc Natl Acad Sci USA. 2015 Apr 7;112(14):4263–7.

46. Piperno DR. 2006. Quaternary environmental history and agricultural impact on vegetation in Central America. Annals MO Botanical Garden 93: 274–296.

47. Emerson BC, Alvarado-Serrano DF, Hickerson MJ. 2015. Model misspecification confounds the estimation of rates and exaggerates their time dependency. Mol Ecol. 24:6013–20.

48. Engelthaler DM, Hicks ND, Gillece JD, Roe CC, Schupp JM, Driebe EM, Gilgado F, Carriconde F, Trilles L, Firacative C, Ngamskulrungroj P, Castañeda E, dos Santos Lazera M, Melhem MSC, Pérez-Bercoff Å, Sorrell TC, Voelz K, May RC, Fisher MC, Thompson GR, Lockhart SR, Keim P, Meyer W. 2014. *Cryptococcus gattii* in North American Pacific Northwest: whole population genome analysis provides insights into species evolution and dispersal. mBio 5:e01464–14

49. Kozarewa I, Turner DJ. 2011. 96-plex molecular barcoding for the Illumina Genome Analyzer. Methods Mol Biol 733:279–98.

50. Bankevich A, Nurk S, Antipov D, Gurevich AA, Dvorkin M, Kulikov AS, Lesin VM, Nikolenko SI, Pham S, Prjibelski AD, Pyshkin AV, Sirotkin AV, Vyahhi N, Tesler G, Alekseyev MA, Pevzner PA. 2012. SPAdes: A New Genome Assembly Algorithm and Its Apphcations to Single-Cell Sequencing. J Comput Biol 19:455–477.

51. DePristo MA, Banks E, Pophn R, Garimella KV, Maguire JR, Hartl C, Philippakis AA, del Angel G, Rivas MA, Hanna M, McKenna A, Fennell TJ, Kernytsky AM, Sivachenko AY, Cibulskis K, Gabriel SB, Altshuler D, Daly MJ. 2011. A framework for variation discovery and genotyping using next-generation DN. sequencing data. Nature Genetics 43:491–498.

52. Swofford DL. 2003. PAUP*: Phylogenetic Analysis Using Parsimony (*and Other Methods). Version 4. Sinauer Associates, Sunderland, Massachusetts.

53. Bryant D, Moulton V. 2004. Neighbor-net: an agglomerative method for the construction of phylogenetic networks. Mol Biol Evol 21:255–65.

54. Huson DH, Bryant D. 2006. Application of phylogenetic networks in evolutionary studies. Mol Biol Evol 23:254–67.

55. Nguyen LT, Schmidt HA, von Haeseler A, Minh BQ. 2015. IQ-TREE: a fast and effective stochastic algorithm for estimating maximum-likehhood phylogenies. Mol Biol Evol 32:268–74.

56. Minh BQ, Nguyen MA, von Haeseler A. 2013. Ultrafast approximation for phylogenetic bootstrap. Mol Biol Evol 30:1188–95.

57. Drummond AJ, Suchard MA, Xie D, Rambaut A. 2012. Bayesian phylogenetics with BEAUti and the BEAST 1.7. J Mol Biol Evol 29:1969–73.

58. Ramasamy RK, Ramasamy S, Bindroo BB, Naik VG. 2014. STRUCTURE PLOT: a program for drawing elegant STRUCTURE bar plots in user friendly interface. Springerplus 3:431.

59. Peakall R, Smouse PE. 2012. GenAlEx 6.5: genetic analysis in Excel. Population genetic software for teaching and research-an update. Bioinformatics 28:2537–2539.

60. Kumar S, Tamura K, Nei M. 1994. MEGA: Molecular Evolutionary Genetics Analysis software for microcomputers. Comput Appl Biosci 10:189–191.

61. Gelman A, Meng X-L. 1998. Simulating normalizing constants: from importance sampling to bridge sampling to path sampling. Statist Sci 13:163–185.

62. Xie W, Lewis PO, Fan Y, Kuo L, Chen MH. 2011. Improving marginal likelihood estimation for Bayesian phylogenetic model selection. Syst Biol 60:150–160.

63. Baele G, Lemey P, Bedford T, Rambaut A, Suchard MA Alekseyenko AV. 2012. Improving the accuracy of demographic and molecular clock model comparison while accommodating phylogenetic uncertainty. Mol Biol Evol 29:2157–2167.

64. Baele G, Li WLS, Drummond AJ, Suchard MA Lemey P. 2013. Accurate model selection of relaxed molecular clocks in Bayesian phylogenetics. Mol Biol Evol. 30:239–243.

65. Rambaut A, Suchard MA, Xie D, Drummond AJ. 2014. Tracer v1.6, Available from http://beast.bio.ed.ac.uk/Tracer

66. Fraser JA, Stajich JE, Tarcha EJ, Cole GT, Inglis DO, Sil A, Heitman J. 2007. Evolution of the mating type locus: insights gained from the dimorphic primary fungal pathogens Histoplasma capsulatum, Coccidioides immitis, and Coccidioides posadasii. Eukaryot Cell 6:622–9.

67. Altschul SF, Gish W, Miller W, Myers EW, Lipman DJ. 1990. Basic local alignment search tooLJMol Biol 215:403–410.

